# Practical probabilistic and graphical formulations of long-read polyploid haplotype phasing

**DOI:** 10.1101/2020.11.06.371799

**Authors:** Jim Shaw, Yun William Yu

## Abstract

Resolving haplotypes in polyploid genomes using phase information from sequencing reads is an important and challenging problem. We introduce two new mathematical formulations of polyploid haplotype phasing: (1) the min-sum max tree partition (MSMTP) problem, which is a more flexible graphical metric compared to the standard minimum error correction (MEC) model in the polyploid setting, and (2) the uniform probabilistic error minimization (UPEM) model, which is a probabilistic analogue of the MEC model. We incorporate both formulations into a long-read based polyploid haplotype phasing method called *flopp*. We show that flopp compares favorably to state-of-the-art algorithms—up to 30 times faster with 2 times fewer switch errors on 6x ploidy simulated data.

## 1 Introduction

As genomic sequencing technologies continue to improve, we are increasingly able to resolve ever finer genomic details. Case in point, traditional genotyping only determines if a particular allele is present in a genome [32]. However, when organisms are *polyploid* (and most eukaryotic organisms are), they have multiple copies of each chromosome. We are then additionally interested in the problem of resolving *haplotypes*, i.e. determining the sequence of alleles on each specific chromosome and not just the presence of an allele within the genome. *Phasing* is the procedure of resolving the haplotypes by linking alleles within a chromosome [7].

We focus on phasing polyploid organisms using third-generation sequencing data. Many plants have *ploidy* greater than two (i.e. have more than two copies of each chromosome), such as tetraploid potatoes (Solanum tuberosum) or hexaploid wheat and cotton. Haplotype phasing has been used to gain insights into evolution [13], breeding [30], and genome-wide association studies [22], among other applications.

The most common way of determining haplotypes is to use pooled genotyping information from a population to estimate haplotypes [7]. For unrelated individuals, sophisticated statistical methods are used to determine the most likely haplotypes for each individual [8,17,12] in the population. For related individuals, identity-by-descent information can be used for haplotype phasing [14, 27]. However, these types of methods do not work on single individuals because they rely on having population data available.

Instead, in this work, we adopt the approach of single individual phasing by sequencing, which is now common in phasing human haplotypes [9]. We focus on using sequencing information for phasing, which allows us to phase a single individual without population information or prior haplotype knowledge. This is closely related to genome assembly where overlapping reads are stitched together [28]; in our case, nearby heterozygous alleles are stitched together by read information. For the rest of the paper, we use the term phasing to mean single individual phasing using sequencing information.

### 1.1 Related work

The first method for phasing polyploid genomes was HapCompass [2], which uses a graphical approach. Popular methods that followed include HapTree [4,5], H-PoP [36] and SDhaP [11]. HapTree and H-PoP heuristically maximize a likelihood function and an objective function based on the MEC model respectively while SDhaP takes a semi-definite programming approach. HapTree-X [5] additionally incorporates long-range expression correlations to allow phasing even of pairs of variants that cannot be covered by a single read, overcoming some of the problems with short-read phasing.

Due to the increased prevalence of long-read data from Oxford Nanopore or PacBio, newer methods taking advantage of the longer-range correlations accessible through long-read data have been proposed [1,33]. Unfortunately, because the error profiles of long-read technologies differ considerably from Illumina short-reads (e.g. a higher prevalence of indel errors compared to SNPs), methods tailored to short-reads such as [25,34] may be ineffective, so altogether new paradigms are required.

At a more theoretical level, in the diploid setting, the standard minimum error correction (MEC) model [6] has proven to be quite powerful. It is known to be APX-Hard and NP-Hard but heuristically solved in practice with relatively good results. Unfortunately, a good MEC score may not imply a good phasing when errors are present [21]. This shortcoming is further exacerbated in the polyploid setting because similar haplotypes may be clustered together since the MEC model does not consider coverage; this phenomenon is known as genome collapsing [33]. Thus, although the MEC model can be applied to the polyploid setting, it may be suboptimal; however, there is yet to be an alternative commonly agreed upon formulation of the polyploid phasing problem. Indeed, this is reflected in the literature: the mathematical underpinnings of the various polyploid phasing algorithms are very diverse.

### 1.2 Contributions

In this paper, we first address the theoretical shortcomings highlighted above by giving two new mathematical formulations of polyploid phasing. We adopt a probabilistic framework that allows us to (1) give a better notion of haplotype similarity between reads and (2) define a new objective function, the uniform probabilistic error minimization (UPEM) score. Furthermore, we introduce the idea of framing the polyploid phasing problem as one of partitioning a graph to minimize the sum of the max spanning trees within each cluster, which we show is related to the MEC formulation in a specific case.

We argue that these formulations are better suited for polyploid haplotype phasing using long-reads. In addition to our theoretical justifications, we also implemented a method we call flopp (fast local polyploid phaser). flopp optimizes the UPEM score and builds up local haplotypes through the graph partitioning procedure described. When tested on simulated data sets, flopp produces much more accurate local haplotype blocks than other methods and also frequently produces the most accurate global phasing. flopp’s runtime is additionally comparable to, and often much faster than, its competitors.

The code for flopp is available at https://github.com/bluenote-1577/flopp. flopp utilizes Rust-Bio [19], is written entirely in the rust programming language and is fully parallelizable. flopp takes as input either BAM + VCF files, or the same fragment file format used by AltHap [16] and H-PoP.

## 2 Methods

### 2.1 Definitions

We represent every read as a sequence of variants (rather than as a string of nucleotides, which is commonly used for mapping/assembly tasks). Let *R* be the set of all reads that align to a chromosome and *m* be the number of variants in our chromosome. Assuming that tetra-allelic SNPs are allowed, every read *r*_*i*_ is considered as an element of the space *r*_*i*_ *∈ {−*, 0, 1, 2, 3*}*^*m*^. A read in this variant space is sometimes called a fragment. Denoting *r*_*i*_[*j*] as the *j*th coordinate of *r*_*i*_, *r*_*i*_[*j*] *∈ {*0, 1, 2, 3*}* if the *j* th variant is contained in the read *r*_*i*_ where 0 represents the reference allele, 1 represents the first alternative allele, and so forth. *r*_*i*_[*j*] = *−* if *r*_*i*_ does not contain the *j* th allele.

We note that flopp by default only uses SNP information, but the user may generate their own fragments, permitting indels and other types of variants to be used even if there are more than four possible alleles. The formalism is the same regardless of the types of variants used or the number of alternative alleles.

For any two reads *r*_1_, *r*_2_, let

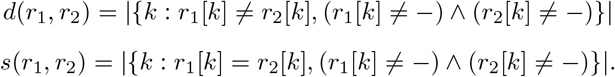

*d* and *s* stand for *different* and *same*, representing the number of different and same variants respectively between two reads.

We use *k* to denote the ploidy. Given a *k*-ploid organism, a natural approach to phasing is to partition *R* into *k* subsets where the cluster membership of a read represents which haplotype the read belongs to. Let *R*_1_, …, *R*_*k*_ be a partition of *R*. Given a partition *P* = *{R*_1_, …, *R*_*k*_*}*, we denote *P* [*i*] = *R*_*i*_.

Define the *consensus haplotype H*(*R*_*i*_) *∈ {−*, 0, 1, 2, 3*}*^*m*^ associated to a subset of reads as follows. For all indices *l* = 1, …, *m* let *H*(*R*_*i*_)[*l*] = arg max*a* |*{r ∈ R*_*i*_ : *r*[*l*] = *a}*| and break ties according to some arbitrary order. If only *−* appear at position *l* over all reads, we take *H*(*R*_*i*_)[*l*] = *−*. It is easy to check that *H*(*R*_*i*_) is a sequence in *{−*, 0, 1, 2, 3*}*^*m*^ such that *H*(*R*_*i*_)[*k*] *≠ −* at indices for which some read overlaps, and 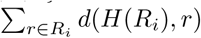 is minimized.

In our formalism, we can phrase the MEC model of haplotype phasing as the task of finding a partition *{R*_1_, …, *R*_*k*_*}* of *R* such that 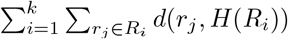, which is called the MEC score, is minimized. For notational purposes, for a sub-set *R*_*i*_ *⊂ R*, define 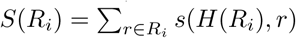 and 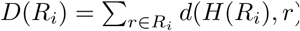. 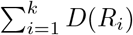 is just the MEC score for a particular partition.

### 2.2 Problem formulation

#### Min-sum max tree partition (MSMTP) model

Let *G*(*R*) = (*R, ϵ, w*) be an undirected graph where the vertices are *R* and edges *E* are present between two reads *r*_1_, *r*_2_ if *r*_1_, *r*_2_ overlap, i.e. *d*(*r*_1_, *r*_2_) + *s*(*r*_1_, *r*_2_) *>* 0. Let the weight of *e* = (*r*_1_, *r*_2_) be *w*(*e*) = *w*(*r*_1_, *r*_2_) for some weight function *w*. We call *G*(*R*) the *read-graph*, see Figure 1. A similar notion is found in [11,1,33,24,23].

**Fig. 1.**
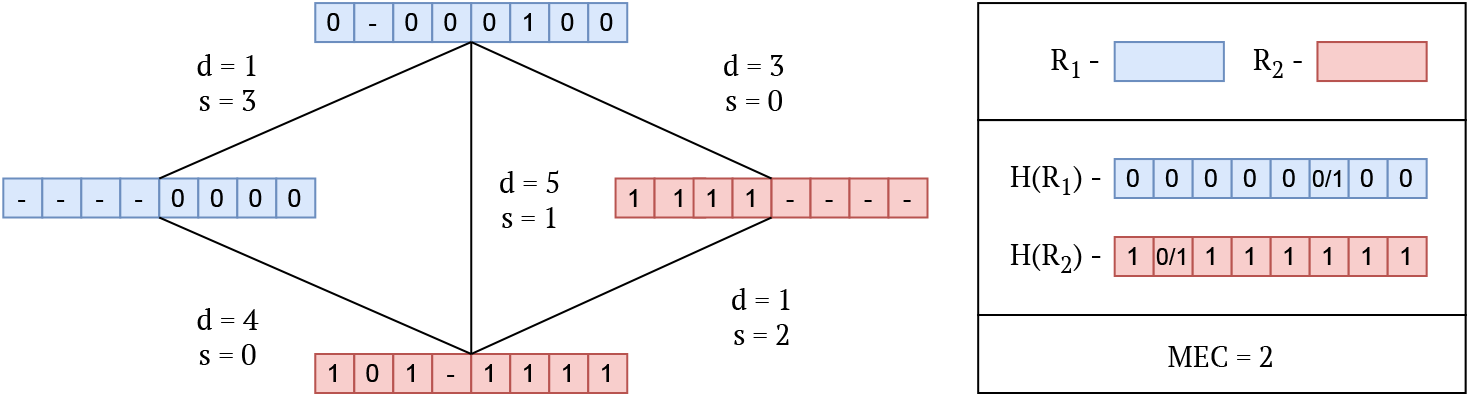
An example of a read-graph (without the weighting *w* specified) along with corresponding *d, s* values between reads. The colors represent a partition of the readgraph into two subsets *R*_1_, *R*_2_. The consensus haplotypes *H*(*R*_1_), *H*(*R*_2_) are shown where the 0*/*1 indicates that either 1 or 0 are valid alleles for a consensus haplotype.

For a partition of *R* into disjoint subsets *{R*_1_, …, *R*_*k*_*}* we take *G*(*R*_*i*_) as defined above. We only consider partitions of vertices such that all *G*(*R*_*i*_) are connected, which we will denote as valid partitions. Let *MST* (*G*) be the maximum spanning tree of a graph *G*. Define

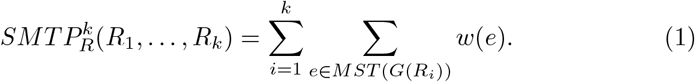

We formulate the *min-sum max tree partition (MSMTP) problem* as finding a valid partition *{R*_1_, …, *R*_*k*_*}* of *R* such that 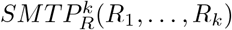 is minimized. We refer to the *objective function being minimized* as the SMTP score and *the computational problem of minimizing the SMTP score* as the MSMTP problem.

The MSMTP problem falls under a class of problems called graph tree partition problems [10], most of which are NP-Hard. We give a proof that MSMTP is NP-Hard in Appendix A.

Intuitively, assuming each *G*(*R*_*i*_) is connected, a maximum spanning tree is a maximum measure of discordance along the entire haplotype. We prove below that under a specific constraint on the read-graph, the SMTP score for *w*(*r*_1_, *r*_2_) = *d*(*r*_1_, *r*_2_) is an upper bound for the MEC score.

##### Theorem 1.

*Suppose w*(*r*_1_, *r*_2_) = *d*(*r*_1_, *r*_2_). *Let a, b ∈* N *and let R be a set of fragments such that for every r ∈ R, for all k ∈ {a, a* + 1, …, *b}, r*[*k*] *≠ − and for l /∈ {a, a* + 1, *…, b}, r*[*l*] = *−. For any R*_*i*_ *in a valid partition {R*_1_, …, *R*_*k*_*} of R*,

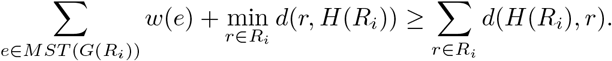

*Therefore*,

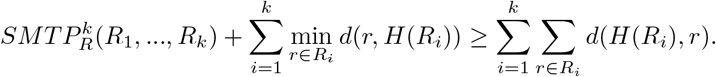

##### Proof.

Take the augmented graph *G*(*R*_*i*_*∪{H*(*R*_*i*_)*}*). It is clear that 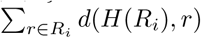 is just 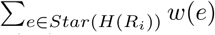, where *Star*(*H*(*R*_*i*_)) is the star-graph having internal node *H*(*R*_*i*_) and every *r ∈ R*_*i*_ as a leaf node.

Now note that *H*(*R*_*i*_) is constructed *precisely* as a sequence in *{−*, 0, 1, 2, 3*}*^*m*^ which is non *−* at indices *a, a*+1, …, *b* that minimizes the sum 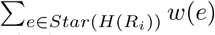. For any *r ∈ R*_*i*_, *r* is also non *−* at the same indices by assumption, so 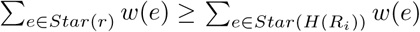.

Removing the node *H*(*R*_*i*_) from *Star*(*r*) and the corresponding edge, we get a spanning tree of *G*(*R*_i_); call this new graph *Star*(*r*)^*i*^. Thus 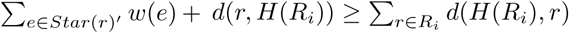, and clearly 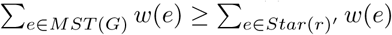. The inequality holds for any *r*, so we can choose *r* to minimize *w*(*r, H*(*R*_*i*_)), finishing the proof.

The above theorem relies on the assumption that all reads in the set overlap exactly. While this is obviously not true for the entire genome, flopp takes a local clustering approach where the entire read set *R* is partitioned into smaller local read sets that overlap significantly.

We verified experimentally that the SMTP score for partitions generated by our method has a strongly linear dependence on the MEC score when *w* = *d*, see Appendix E. These results justify that minimizing the SMTP score is worthwhile for this specific case. However, we do not have to necessarily use *w* = *d*. In Section 2.3 and we opt for a more theoretically sound probabilistic weighting.

#### Uniform probabilistic error minimization (UPEM) model

The SMTP score has problems with collapsing genomes in the same manner the MEC score does; it does not take into account the assumption that coverage should be uniform between haplotypes. Concretely, if a polyploid organism has two identical haplotypes, the reads from both haplotypes may be clustered together in the MEC model and a noisy read may instead be placed in its own cluster.

Let *ϵ* represent the probability that a variant is called incorrecly. Let *σ ∈* ℝ be a normalizing constant, and *X*_*i*_ *∼Binomial* (⌈*D*(*R*_*i*_) + *S*(*R*_*i*_) */σ* ⌉, *ϵ*) be a binomial random variable. Then

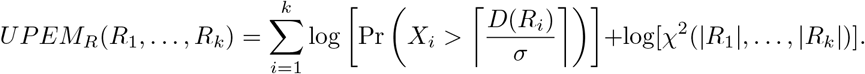

The *χ*^2^(*x*_1_, …, *x*_*n*_) term is the p-value for the *χ*^2^ test while the binomial term is a sum of log one-sided binomial tests where the null hypothesis is that the error rate of a clustering is *ϵ*. Therefore the UPEM score is just a sum of log p-values.

The UPEM score is a probabilistic version of the MEC model under the hypothesis that the errors and coverage are uniform across haplotypes. The parameter *σ* is a normalizing constant and is important because if a specific genome has a high rate of heterozygosity and *ϵ* is slightly underestimated, then the sample size *D*(*R*_*i*_) + *S*(*R*_*i*_) is large. The p-value associated with the binomial random variable will be extremely small and drown out the contributions from the *χ*^2^ term, so *σ* is a learned data set specific constant used to keep the two terms balanced.

UPEM maximization enforces a relatively uniform partition. Furthermore, errors will be distributed among partitions equally due to the non-linearity of the UPEM score; if one cluster is extremely erroneous, the sum of the binomial terms may be higher for a more spread out error even if the overall MEC score is slightly higher. If error and coverage uniformity assumptions are satisfied, clearly these two properties are desirable in an objective function.

### 2.3 Local graph clustering

We now discuss the algorithms implemented in flopp. A high-level block diagram showing outlining flopp’s main processing stages are outlined in Figure 2. Importantly, flopp is a local clustering method, which means that we first attempt to find good partitions for smaller subsets of reads by optimizing the SMTP and UPEM functions and then joining haplotype blocks together afterwards.

**Fig. 2.**
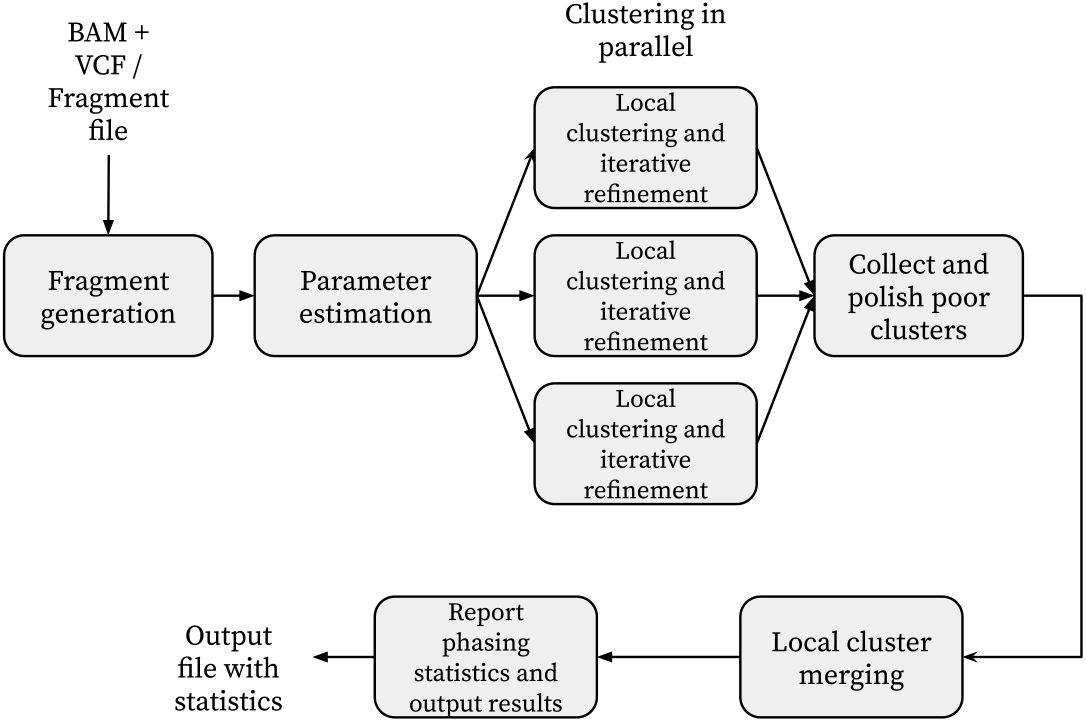
A block diagram outlining flopp’s major processing steps. flopp takes BAM+VCF files or a fragment file as input. The fragments are clustered into local haplotype blocks, which are then polished and merged. The final phasing is output to the user as well as various phasing statistics.

#### Choice of edge weight function

Previous methods that use a read-graph formalism such as SDhaP [11], FastHap [24], WHP [33], and nPhase [1] all define different weightings between reads. Two previously used weight functions are

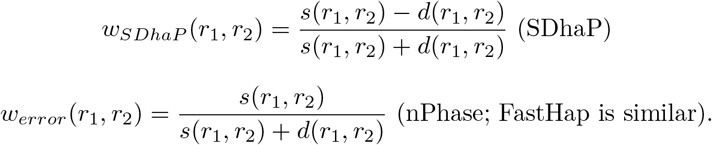

These weightings are quite simple and have issues when the length of the reads have high variance, as is the case with long-reads. A more sophisticated approach is to use a probabilistic model. Let *E* (*r*_1_, *r*_2_) = 1 *− w*_*error*_(*r*_1_, *r*_2_) be the average error rate between two reads. Assuming an error parameter *ϵ* as described in the UPEM formula, we define

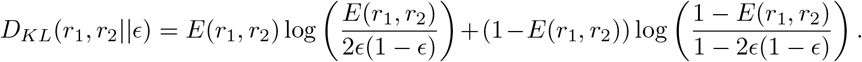

*D*_*KL*_(*r*_1_, *r*_2_ *∥ ϵ*) is the Kullback-Leibler divergence between a *E*(*r*_1_, *r*_2_) *—* coin and a 2*ϵ*(1*− ϵ*) coin. The reason we use 2*ϵ*(1*−ϵ*) is because, given two reads with error rate *ϵ*, the probability that two variants from the same haplotype are different is 2*ϵ*(1 *− ϵ*). Now let

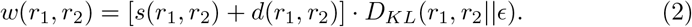

*D*_*KL*_(*r*_1_, *r*_2_ *ϵ*) is a measure of divergence between the error profiles of the two reads and the expected error profile for two reads from the same haplotype. There is a slight issue that *D*_*KL*_(*r*_1_, *r*_2_*∥E*) has a minimum at *E* (*r*_1_, *r*_2_) = 2*ϵ*(1*−ϵ*) when *ϵ* is fixed. We want *D*_*KL*_ to be a monotonically increasing function with respect to *E*, so we flip the sign of *D*_*KL*_(*r*_1_, *r*_2_*∥ϵ*) if *E <* 2*ϵ*(1 *− ϵ*).

There is a nice interpretation of *w*(*r*_1_, *r*_2_) when *E* (*r*_1_, *r*_2_) *>* 2*ϵ*(1 *ϵ*) as given by the following theorem.

##### Theorem 2 (Arratia and Gordon [3])

*Suppose p < a <* 1. *Let* 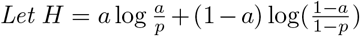 *be the Kullback-Leibler divergence between an a−coin and a p−coin. Then*

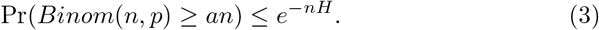

*In particular, this bound is asymptotically tight up to logarithms, i*.*e*.

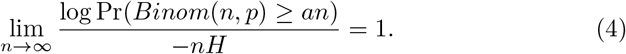

The proof of the above theorem is simple; it follows by an application of a Chernoff bound and Sterling’s inequality. In [3], they show that this approximation is particularly good when *a ≫ p*. Theorem 2 gives an interpretation of *w*(*r*_1_, *r*_2_) as the negative log value of a 1-sided binomial test where the null hypothesis is that *r*_1_, *r*_2_ come from the same haplotype, the number of trials is *s*(*r*_1_, *r*_2_) + *d*(*r*_1_, *r*_2_) and the number of successes is *d*(*r*_1_, *r*_2_).

WHP [33] uses a log ratio of binomial probability densities as their edge weight. Mathematically, the resulting formula is almost the same as Equation 2, except they fix *E*(*r*_1_, *r*_2_) to be the average error rate between reads from different haplotypes, a constant. They end up with both negative and positive edges with large magnitude and Theorem 2 does *not* apply to their case. On the other hand, our edge weights are rigorously justified as negative log p-values for a binomial test, making it more interpretable.

#### MSMTP Algorithm

We devised a heuristic algorithm for local clustering inspired by the MSMTP problem. From now on, let *S ⊂ R*. The pseudocode is shown in Algorithm 1. In the diploid setting, the main idea behind this algorithm is similar to that of FastHap [24], but the algorithm is quite different after generalizing to higher ploidies.

##### Algorithm 1 Greedy min-max read partitioning

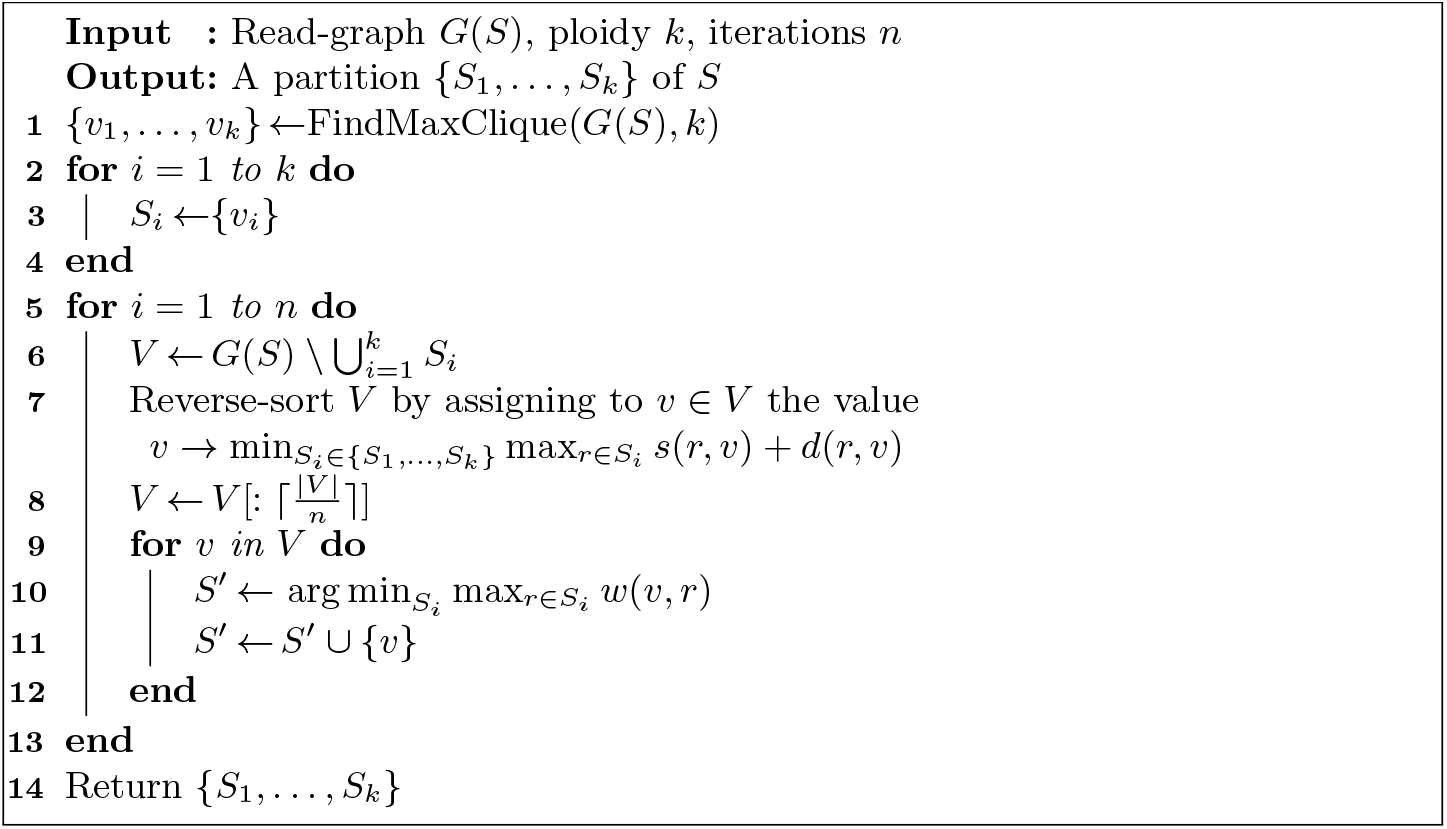

For the FindMaxClique method mentioned in Algorithm 1, we use a greedy clique-finding method which takes *O*(*k*|*S*| log |*S*| + *k*^2^|*S*|) time. First, we take the heaviest edge between two vertices and make this our 2-clique. Then we re-sort vertices by maximizing the minimum distance from the vertex to all vertices in the 2-clique, add a third edge to get a 3-clique, and so forth until we get a *k−*clique.

The complexity of the local clustering algorithm is *O*(*n* |*S*| ^2^ + *k* |*S*| log |*S*| + *k*^2^ |*S*|). In practice, note that |*S*| *≫k*. The parameter *n* is fixed to be 10. By iterating over *n*, we re-sort the edges based on their overlaps to the new clusters, which have changed since the previous iteration. This improves the order in which we add vertices to the clusters.

The connection to MSMTP is at line 10. A priori, it is not obvious what metric to use to determine which cluster to put the vertex in. For Kruskal’s algorithm, one starts by considering the heaviest edges, so we decided to minimize the maximum edge from the vertex to the cluster so that the heaviest edges are small. Intuitively, a maximum spanning tree is sensitive to a highly erroneous edge, so we prioritize minimizing the worst edge even if on average the vertex connects to the cluster well.

#### Iterative refinement of local clusters

We refine the output of the local clustering procedure by optimizing the UPEM score using a method similar to the Kernighan-Lin algorithm. Pseudocode can be found in the appendix as Algorithm 2.

The algorithm takes in a partition *P* = *{S*_1_, …, *S*_*k*_ *}* of a subset of reads *S ⊂R* as well as a parameter *n* which indicates the maximum number of iterations. The algorithm checks how moving reads from one partition to another partition changes the UPEM score for every read. We store the best moves and execute a fraction of them. Proceed for *n* iterations or until the UPEM score does not improve anymore. In practice, we set the parameter *n* = 10. The time complexity of Algorithm is *O*(*n*|*S*|*k* log(|*S*|*k*)).

#### Local phasing procedure

Note that Algorithms 1 and 2 work on subsets of reads or subgraphs of the underlying read-graph. Let *b∈* N be a constant representing the length of a local block. We consider subsets *B*_1_, …, *B*_*l*_ *⊂ R* where

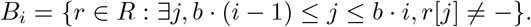

The subsets are just all reads that overlap a shifted interval of size *b*, similar to the work done in. After choosing a suitable *b*, we run the read-partitioning and iterative refinement on all *B*_1_, …, *B*_*l*_ to generate a set of partitions *P*_1_, …, *P*_*l*_. We found that a suitable value of *b* is the 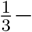 quantile value of read lengths. By read length we mean the last non ^*′*^*−*^*′*^ position minus the first non ^*′*^*−*^*′*^ position of *r ∈ {*0, 1, 2, 3, *−}*.

It is important to note that computationally, the local clustering procedure is easily parallelizable. The local clustering step has therefore a linear speed-up in the number of threads.

### 2.4 Polishing, merging, and parameter estimation

#### Filling in erroneous blocks

Once we obtain a set of partitions *P*_1_, …, *P*_*l*_ of *B*_1_, …, *B*_*l*_ according to the above local clustering procedure, we can identify partitions with low UPEM score and correct them. We describe this procedure in Appendix C.

#### Local cluster merging

Let *P* represent the final partition of all reads *R*. We build *P* given *P*_1_, …, *P*_*l*_ as follows. Start with *P* = *P*_1_. Let *S*_*k*_ be the symmetric group on *k−* elements, i.e. the set of all permutations on *k* elements. At the *i*th step of the procedure, let

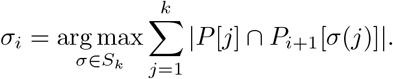

Then let *P* [*j*] = *P* [*j*] *∪P*_*i*+1_[*σ*_*i*_(*j*)] for all *j* = 1, …, *k*. Repeat this procedure for *i* = 1, …, *l −*1.

We experimented with more sophisticated merging procedures such as a beam-search approach but did not observe a significantly better phasing on any of our data sets.

#### Phasing output

Once a final partition *P* has been found, we build the final phased haplotypes. The output is *k* sequences in *{*0, 1, 2, 3, *−}* ^*m*^ where *m* is the number of variants. flopp can take fragment files as input, in which each line of the file describes a single read fragment in *{*0, 1, 2, 3, *−}* ^*m*^, or it can take a VCF file and a BAM file of aligned reads. In the former case, without a VCF file, we do not have genotyping information, so we simply output *{H*_1_, …, *H*_*k*_*}* where *H*_*i*_ = *H*(*P* [*i*]) is the consensus haplotype. If a VCF file is present, flopp constrains the final haplotype by the genotypes, that is, for some output variant, the number of reference and alternate alleles is the same as in the VCF file. We describe this procedure in Appendix.

#### Parameter estimation

We already mentioned in previous sections how we set all parameters for algorithms except for *ϵ* and the parameter *σ* in the UPEM score. We set *σ* to be the median length of the reads divided by 25, which we found empirically to be a good balance between the binomial and chi-squared terms.

To estimate *ϵ*, we start with an initial guess of *ϵ* = 0.03. We then select 10 subsets *B*_*i*_ *⊂ R* at random and perform local clustering and refinement. We estimate *ϵ* from the 10 *S· k* total clusters by choosing *ϵ* = the 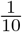 th quantile error, where for each cluster *C ⊂ B*_*i*_ the error is 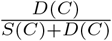. We pick a bottom quantile because we assume that there is some error in our method, so to get the true error rate we must underestimate the computed errors.

## 3 Results and Discussion

### 3.1 Phasing metrics

There are a plethora of phasing metrics developed for diploid and polyploid phasing [26]. We use three different metrics of accuracy, but argue that each individual metric can be misleading and that all three metrics should be used in unison before drawing conclusions on phasing accuracy.

For a global measure of goodness of phasing, we use the Hamming error rate. Given a set of true haplotypes *H* = *{H*[1], …, *H*[*k*]*}* and a set of candidate haplotypes *H*^*\**^ = *{H*^*\**^[1], …, *H*^*\**^[*k*]*}*, we define the Hamming error rate as

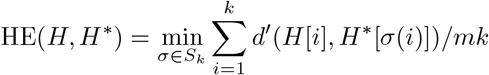

where *S*_*k*_ is the set of permutations on *k* elements, *m* is the length of each *H*[*i*], *k* is the ploidy, and *d*^*′*^ is the same as the *d* function defined before except we count the case where one haplotype has a ^*′*^*−*^*′*^ at a coordinate as an error.

We define the switch error rate (SWER) similarly to WHP [33]. Let *Π*_*i*_ *⊂S*_*k*_ be the set of permutations such that *H*[*j*][*i*] = *H*[*j*][*σ*(*i*)] for all 1*≤ j ≤k*. These are the mappings from the truth to the candidate haplotypes that preserve the alleles at position *i*. Then we define the switch error as

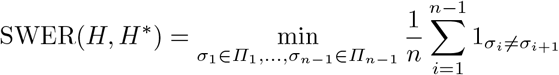

where 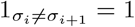 if *σ*_*i ≠*_ *σ*_*i*+1_ and 0 otherwise.

The Hamming error, while easily interpretable, can be unstable. A single switch error can drastically alter the Hamming error rate. The SWER also has issues; for example, if two switches occur consecutively, the phasing is still relatively good but the SWER is worse than if only one switch occurred.

We define a new error rate, called the q-block error rate. For a haplotype *H*[*i*], break *H*[*i*] into non-overlapping substrings of length *q*. Denote each new block 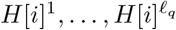. For a set of haplotypes *H*, doing this for every haplotype gives a collection of haplotype blocks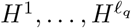. Then the q-block error rate is

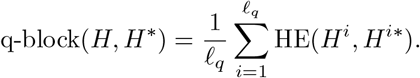

The q-block error rate measures local goodness of assembly and interpolates between the Hamming error rate, when *q* is the size of the genome, and a metric similar to the switch error when *q* = 2.

### 3.2 Simulation procedure

We used the v4.04 assembly of Solanum tuberosum produced by the Potato Genome Sequencing Consortium [15] as a reference. We took the first 3.5 Mb of chromosome 1 and removed all “N”s, leaving about 3.02 Mb and simulated bi-allelic SNPs using the haplogenerator software [26], a script made specifically for realistic simulation of variants in polyploid genomes.

We generated SNPs with mean distance between SNPs of 45 bp. This is in line with the 42.5 bp average distance between variants as seen in [35] for Solanum tuberosum; in that study they observe that *>* 80% of variants are biallelic SNPs. This is also in line with the 58 bp mean distance between variants seen for the hexaploid sweet potato (Ipomoea batatas) observed in [38]. The dosage probabilities provided to haplogenerator are the same parameters as used in [26] for the tetraploid case. When simulating triploid genomes, we use the same dosage probabilities but disallow the case for three alternate alleles. We repeated our test for mean distance between SNPs of 90 and 135 bp in Appendix G.

We used two different software packages for simulating reads. We used PaSS [39] with the provided default error profiles for simulating PacBio RS reads. PaSS has higher error rates than other methods like PBSIM [29] which tends to underestimate error [39]. We used NanoSim [37] for simulating nanopore reads using a default pre-trained error model based on real human dataset provided by the software.

After generating the haplotypes and simulating the reads from the haplotypes, we obtain a truth VCF from the haplotypes. We map the reads using minimap2 [20] to the reference genome. The scripts for the simulation pipeline can be found at https://github.com/bluenote-1577/flopp_test.

### 3.3 Results on simulated data set

We primarily test against H-PoPG [36], the genotype constrained version of H-PoP. We tried testing against WHP, but found that the results for WHP were extremely discontiguous and the accuracy was relatively poor across our data sets. We show an example of this in Appendix F. Other methods such as HapTree and AltHap were tested, but we ran into issues with either computing time or poor accuracy due to the methods not being suited for long-read data. We did not test against nPhase because the output of nPhase does not have a fixed number of haplotypes.

The switch error and Hamming error rates are shown in Figure 3. For each test, we ran the entire pipeline three times; each run at high ploidies takes on the timescale of days to complete. The run times on PacBio reads for H-PoPG, flopp, as well as one instance of WHP on 3x ploidy data are shown in Figure 4.

**Fig. 3.**
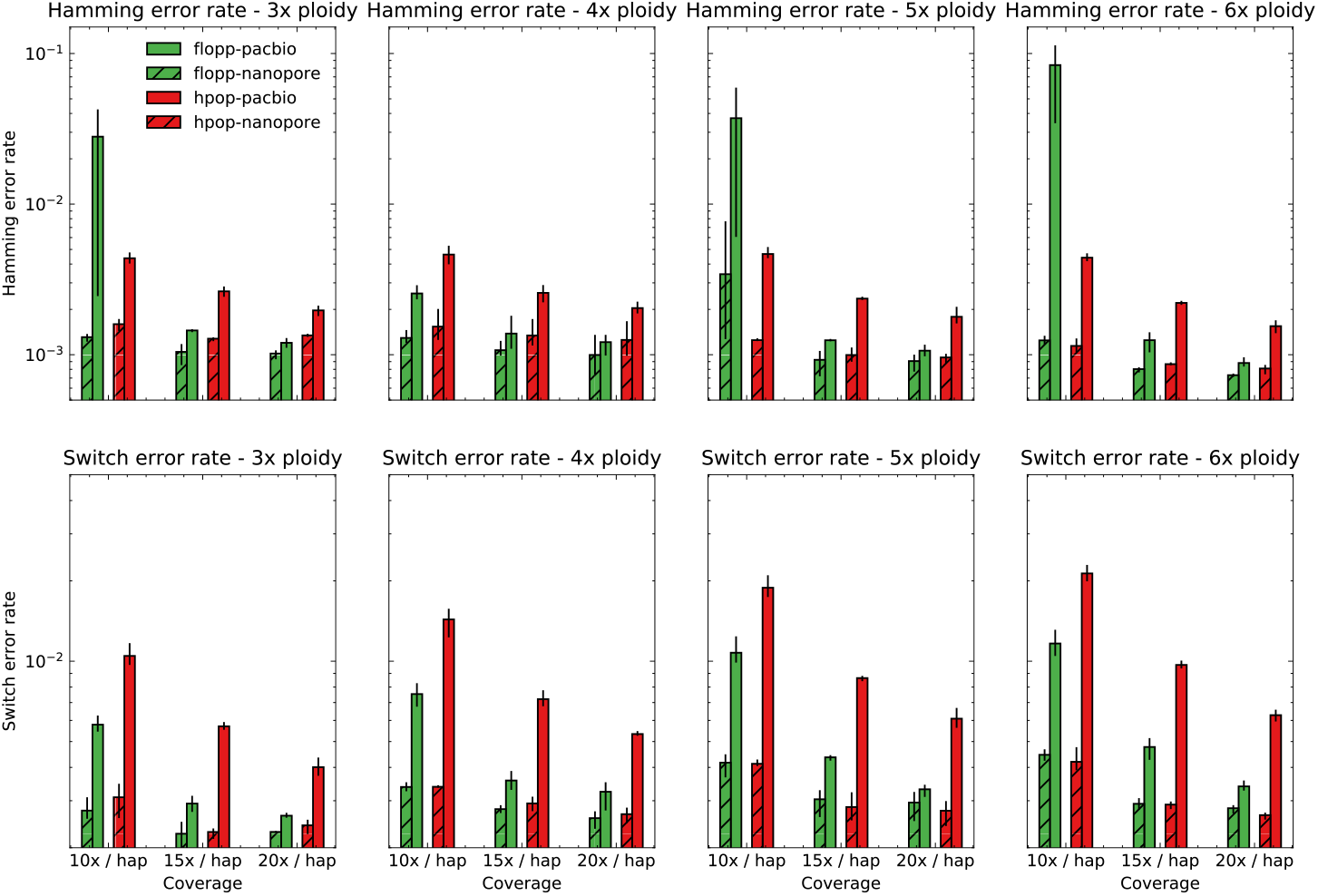
The mean switch error rate and Hamming error rate from testing on simulated data sets as described in Section 3.2 over a range of ploidies and coverages on two different types of read simulators. The error bars represent the lowest and highest values for the metric over three iterations. The results on the Nanopore simulated reads from NanoSim [37] are similar for both methods. flopp achieves much better switch error rates on the PacBio simulated reads from PaSS [39], although for low coverage flopp sometimes has a higher Hamming error rate.

**Fig. 4.**
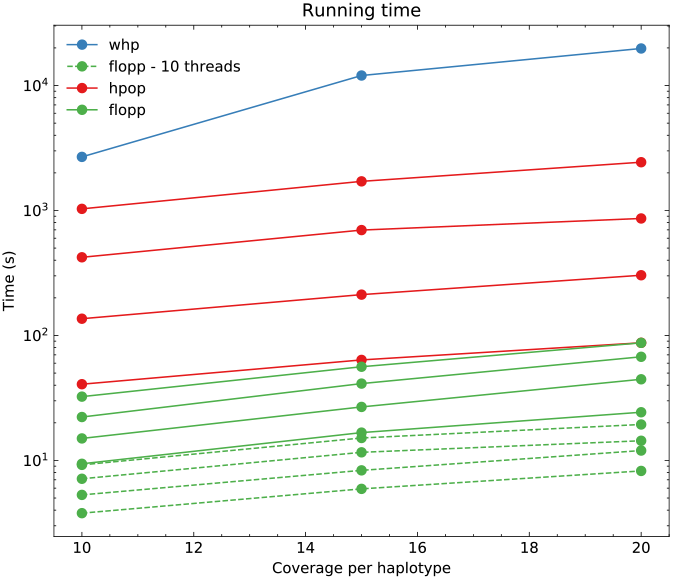
Run times for flopp, H-PoPG, and one instance of WHP. Each line represents a different ploidy with higher ploidies taking longer. The one instance of WHP was run at 3x ploidy.

For the nanopore data set simulated from NanoSim, the results for H-PoPG and flopp are very similar. The PaSS PacBio simulator outputs reads that are more erroneous, and we can see that flopp generally performs better than H-PoPG across ploidies except for the Hamming error rate when the coverage is relatively low; interestingly, the switch error rate is still lower in this case. flopp’s switch error rate is consistently 1.5-2 times lower than H-PoPG for the simulated PacBio reads. On the low coverage data sets, H-PoPG’s global optimization strategy leads to a better global phasing than flopp’s local methodology.

Note that in these tests we phase a contig of length 3.02 Mb, whereas most genes of interest are *<* 50 kb. In Figure 5 we plot the mean q-block error rates of the 5x and 6x ploidy phasings at 10x coverage; these runs have higher Hamming error rates for flopp than H-PoPG. For blocks of up to around 850 · 45 = 38250 bases, flopp outputs phasings with lower q-block error rates than H-PoPG despite a larger global error rate. Although flopp may sometimes give worse global phasings than H-PoPG, flopp can give extremely accurate local phasings.

**Fig. 5.**
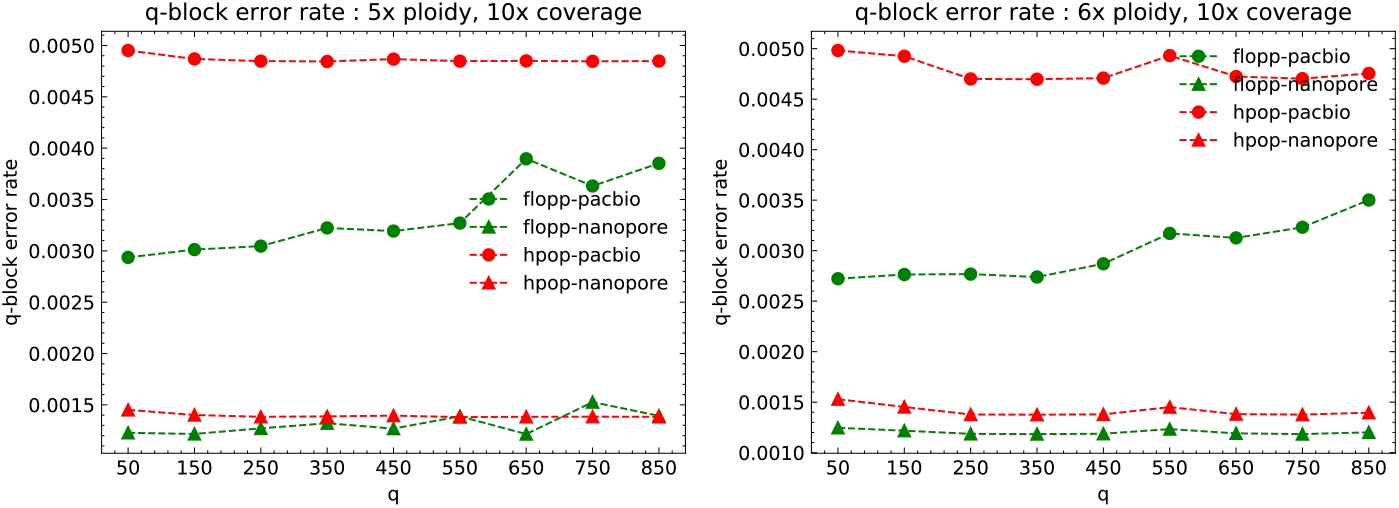
The mean q-block error rates for the 5x and 6x ploidy data sets at 10x coverage per haplotype over three iterations. Each variant is spaced on average 45 bp apart, so each block is of size *∼ q ·* 45. The upward trend indicates an inaccurate global phasing, but the q-block error rates for flopp are lower than for H-PoP in this regime.

Computationally, Figure 4 shows that flopp is at least 3 times faster than H-PoPG and up to 30 times faster than H-PoPG for 6x ploidy on a *single thread*.

The local clustering step sees a linear speedup from multi-threading. For most data sets, we get a 3-4 times speedup with 10 threads.

## 4 Conclusion

In this paper, we presented two new formulations of polyploid phasing, the minsum max tree partition (MSMTP) problem and the uniform probabilistic error minimization (UPEM) model. The SMTP score is a flexible graphical interpretation of haplotype phasing which is related to the MEC score when using a specific weighting on the read-graph, whereas the UPEM score is a superior version of the MEC score when uniformity assumptions are satisfied. Using our probabilistic formulation, we give a new notion of distance between read fragments based on the Kullback-Leibler divergence which has a rigorous interpretation as a log p-value of a one-sided binomial test.

We implemented a fast, local phasing procedure using these new formulations and showed that our software, flopp, is faster and more accurate on high coverage data while always having extremely accurate local phasings across a range of error profiles, coverages, and ploidies.

## Acknowledgments

This work was supported by a faculty startup fund from the University of Toronto at Scarborough Department of Computer and Mathematical Sciences.

## A NP-Hardness of MSMTP

Let *G* = (*V, ϵ, w*) be a connected, undirected, and weighted graph with weight function *w*. Call a partition *V*_1_, …, *V*_*k*_ of the vertices into *k* disjoint sets valid if each *G*(*V*_*i*_) is connected. Then the min-sum max tree partition (MSMTP) problem is to find a valid partition of *V* into *k* sets that minimizes 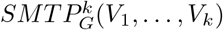.

### Theorem 3.

*MSMTP is NP-Hard for k ≥* 3.

### Proof.

We take a reduction from graph coloring.

Let *G*^*′*^ = (*V,E*^*′*^, *w*^*′*^) be a complete graph where the vertices of *G*^*′*^ are the same as *G*. Let the weight of any edge *e ∈E*^*′*^ to be 2 if *e ∈E*, the original graph, and let it be 1 otherwise.

Let *V*_1_, …, *V*_*k*_ be a (valid) partition of *V* that solves *MSMTP* for *G*^*′*^. Note that none of *V*_1_, …, *V*_*k*_ are empty, otherwise we could move a single vertex to the empty set and it would have a lower *SMTP* value for this graph. Therefore the total number of edges over all spanning trees of *G*^*′*^ (*V*_1_), …, *G*^*′*^ (*V*_*k*_) is 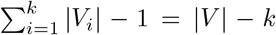. I claim that a k-coloring of *G* exists if and only if 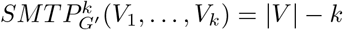.

If 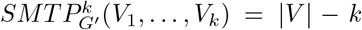, then the maximum spanning tree for each subset only contains edges with weight 1. In particular, this means that no subgraph *G*^*′*^ (*V*_*i*_) has an edge with weight 2, so there are no edges between any vertices of *V*_*i*_ in *G*, otherwise the weight 2 edge would be included in the max spanning tree. Thus the partition *V*_1_, …, *V*_*k*_ gives a *k* coloring of *G*.

For the other implication, clearly if a *k −*coloring exists for *G*, then we can find a partition *V*_1_, …, *V*_*k*_ such that *G*^*′*^(*V*_*i*_) only has edges of weight 1 between vertices. Then 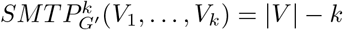follows.

Therefore any algorithm which solves MSMTP also decides if *G* has a k-coloring, which is NP-Complete for *k ≥* 3.

## B Iterative refinement algorithm

We present the pseudocode for the iterative UPEM maximization procedure that follows local clustering in Algorithm 2. In lines 4-15, we check how swapping vertices between partitions affects the overall UPEM score and take a fraction of the best swaps in line 15. We then execute the swaps and check if the UPEM score has increased. If it has not, we terminate the algorithm, otherwise we continue until *n* iterations has passed. We take *n* = 10 in practice, and note that almost always the algorithm terminates before 10 iterations pass.

## C Correcting erroneous haplotype blocks by filling

After obtaining a set of partitions *P*_1_, …, *P*_*l*_ of subsets of reads *B*_1_, …, *B*_*l*_ according to the local haplotyping procedure described in Section 2.3, we correct poor clusters by the following procedure. After computing the UPEM scores for every partition, we use a simple 3.0 IQR (inter-quartile range) outlier detection method for the distribution of UPEM scores. For an outlying partition *P*_*i*_ of the read set *B*_*i*_, if a partition *P*_*i−*1_ is not an outlier, we remove *P*_*i*_ and extend *P*_*i−*1_ to include *B*_*i*_. To do this, we run a subroutine of Algorithm 1 where we skip the clique finding procedure and instead treat the partition *P*_*i−*1_ as the initial clusters. We then run lines 9-12 of Algorithm 1 where in line 9 we iterate over *v ∈ B*_*i*_ *\ B*_*i−*1_ instead.

### Algorithm 2 Iterative refinement of UPEM score

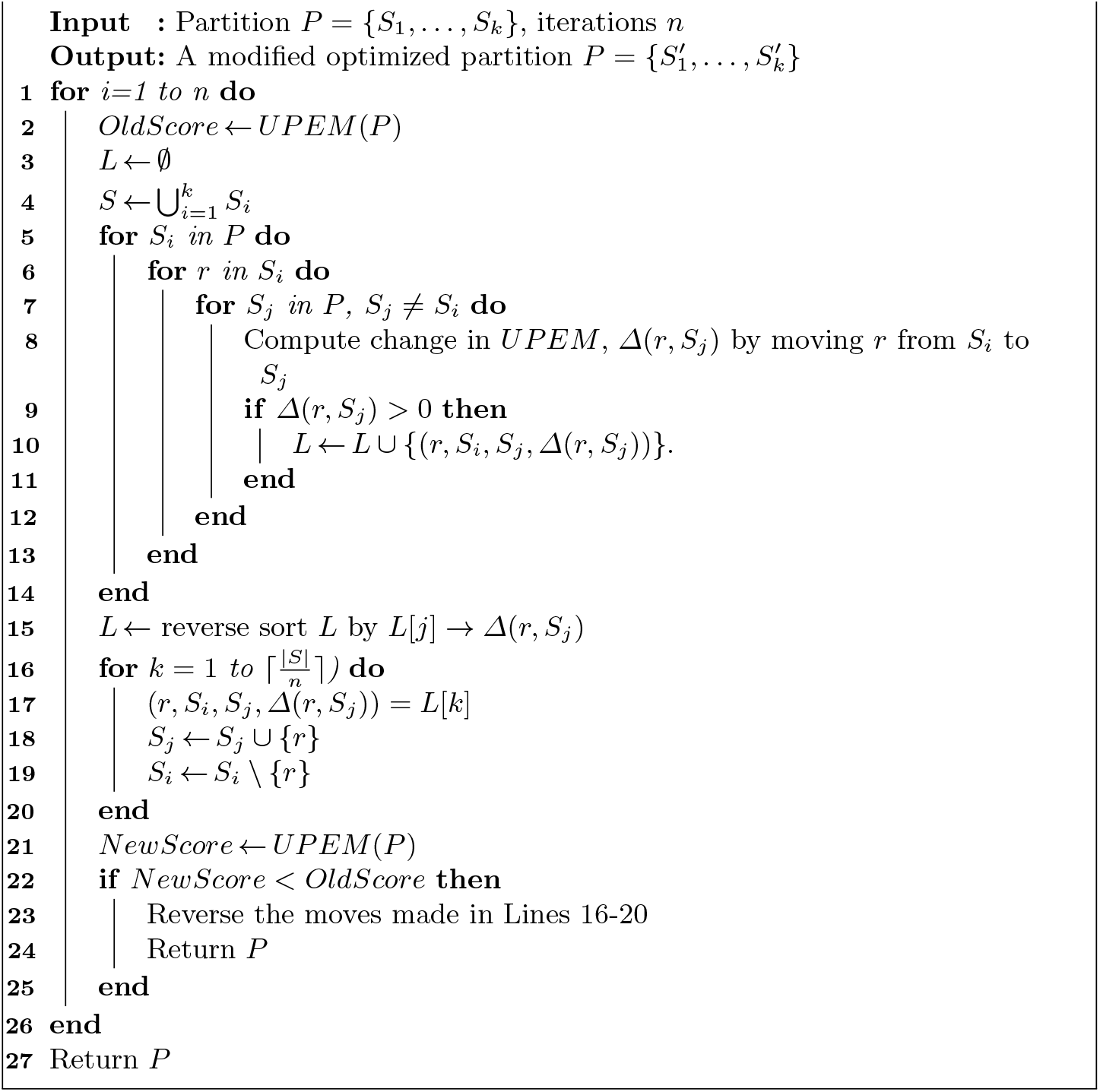

## D Genotype polishing using VCF

We constrain the haplotypes using the VCF as follows. For every variant indexed over 1 *≤ i ≤ m*, let *c*(*i, j, a*) *∈* ℝ be a value representing the confidence of calling allele *a* at index *i* for the haplotype represented by *P* [*j*]. We produce *k−*haplotypes according to Algorithm 3.

### Algorithm 3 Polishing output haplotypes using genotype information.

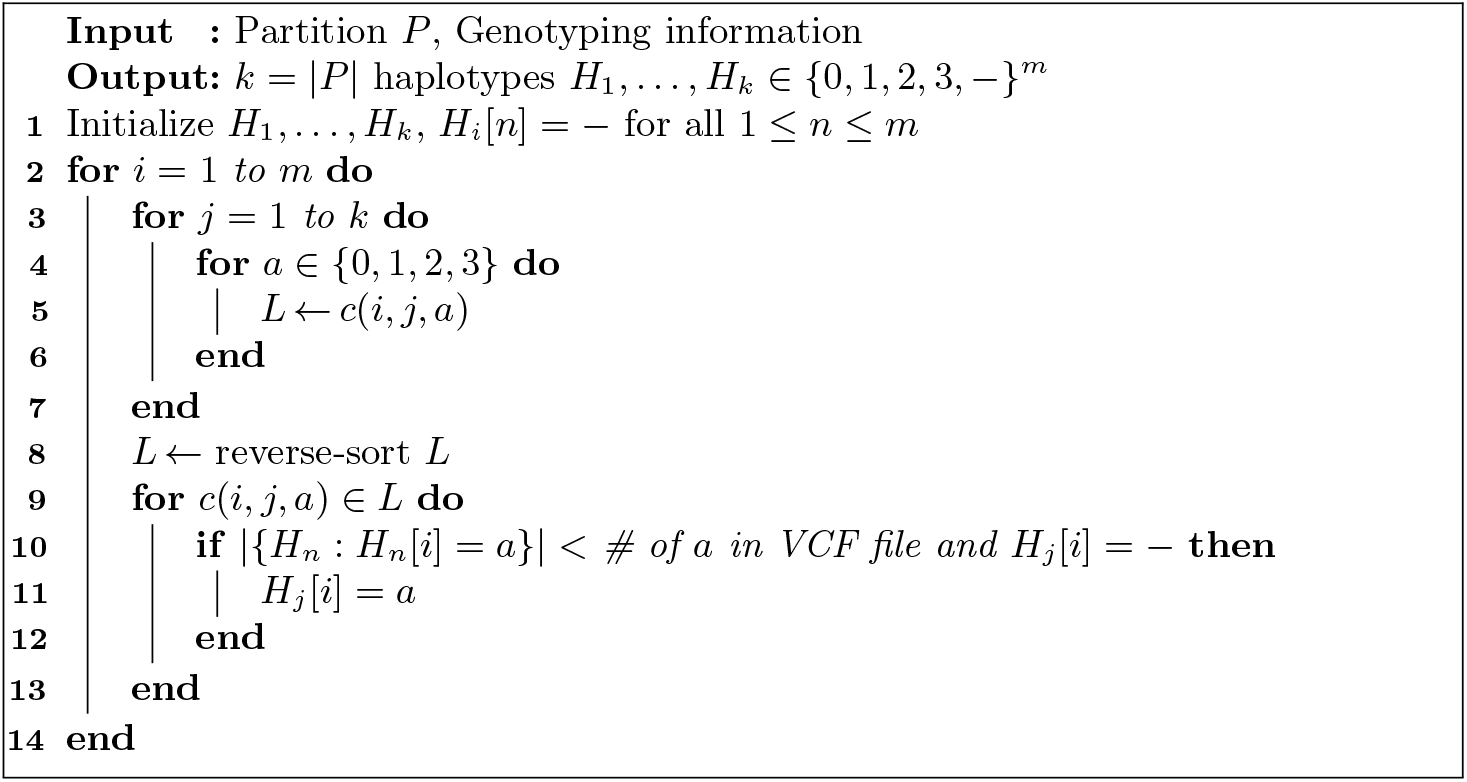

For the function *c*(*i, j, a*) describing the confidence for calling allele *a* at position *m* for haplotype *j*, we choose the function

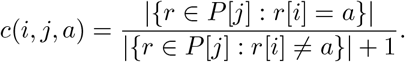

Note that H-PoP uses the same method for polishing, but they choose a confidence function that is a difference instead of a ratio. A ratio does a better job because for a particular haplotype if there are 100 reads that have allele 1 at position 1, and 50 reads that have allele 2, we believe that this is a less confident call than a haplotype with 50 reads with allele 1 and 0 reads that have other alleles.

## E MSMTP vs MEC

We ran flopp on four different simulated datasets (see Section 3.2) and calculated the SMTP score with the weight function *w*(*r*_1_, *r*_2_) = *d*(*r*_1_, *r*_2_), see Equation 1, and the MEC score for each local partition.

We varied the coverage between 10x to 20x for a simulated 4x ploidy genome. We also varied the length of the local partition blocks by manually changing the parameter *b* mentioned at the end of Section 2.3 over three different values for each different data set to investigate how the size of the local clusters affects the SMTP and MEC relationship. The results are shown in Figure 6

**Fig. 6.**
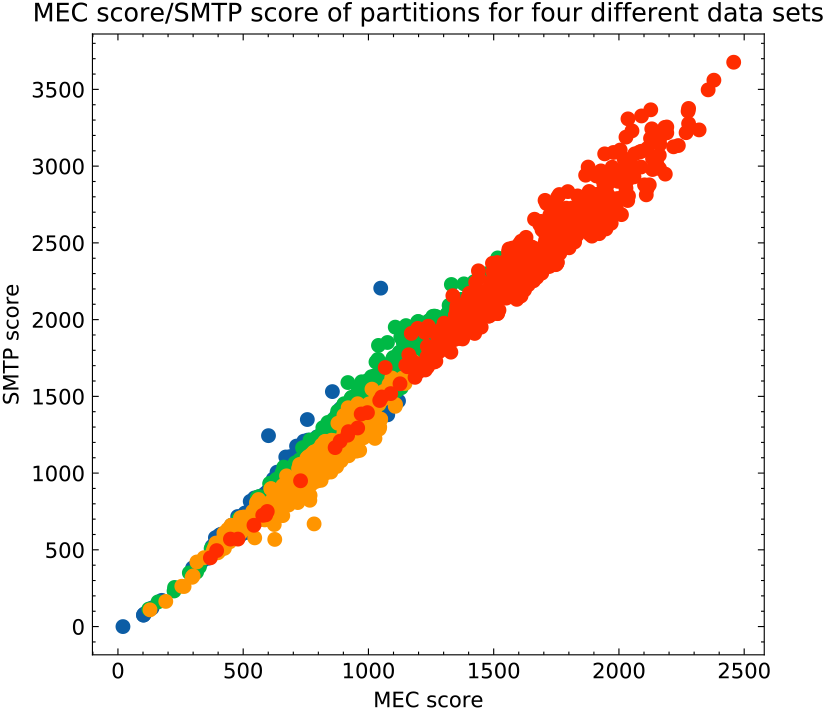
A plot of the SMTP vs MEC score for every local cluster generated by flopp over four different simulated data sets across different coverages with each color representing a distinct data set.

## F WHP Results on 3x ploidy data

We ran WhatsHap Polyphase (WHP) on the simulated genomes as described in Section 3.2. We ran WHP with default settings. In particular, the block-cut sensitivity setting which determines the contiguity of the blocks was set at the default value of 4. There were around 67000 variants in the contig of 3.02 Mb; The N50 over all iterations, coverages, and read types of WHP’s output never exceeded 20. The results are shown in Figure 7.

**Fig. 7.**
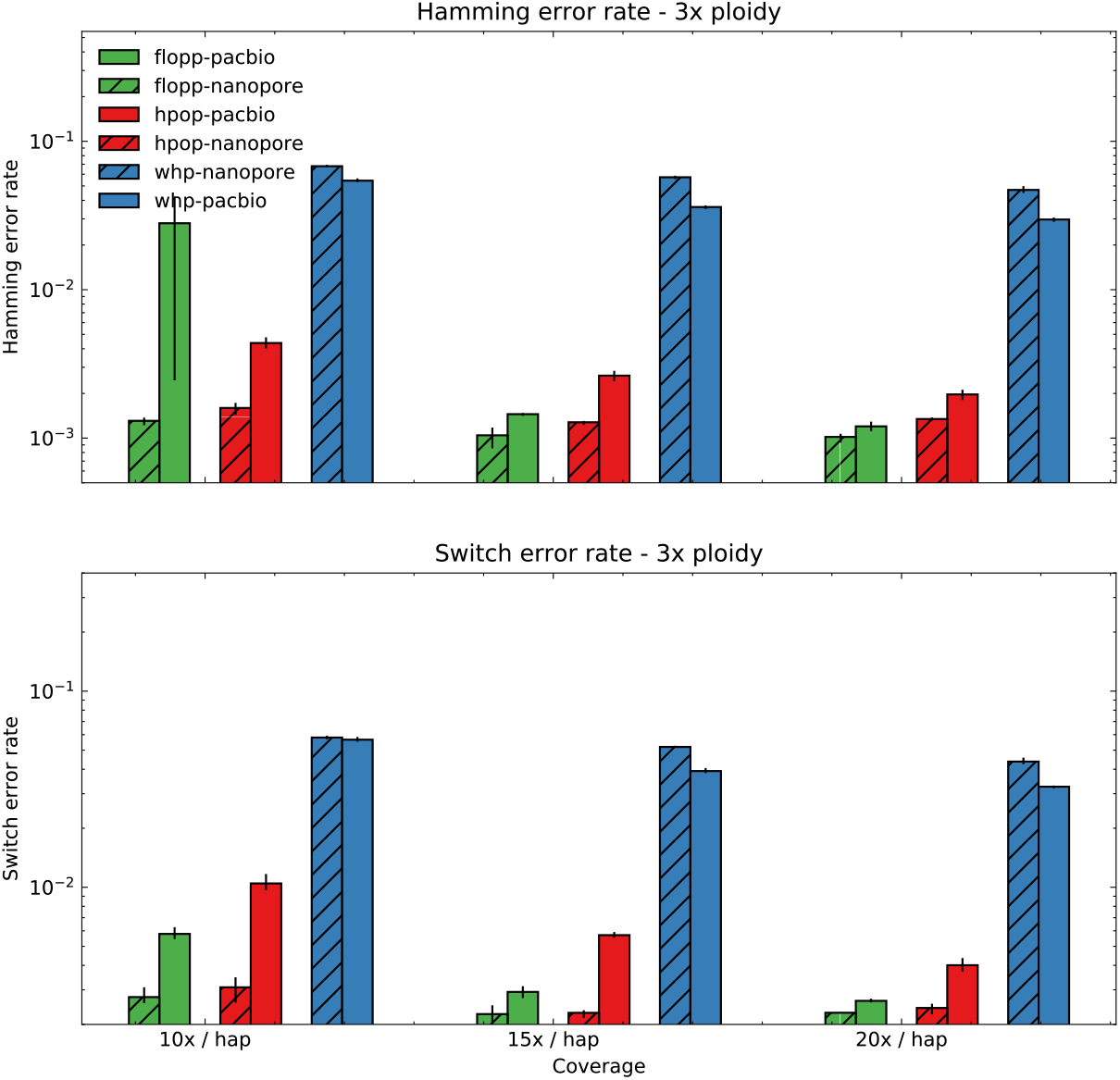
SWER and Hamming error rates of flopp, H-PoPG, and WHP on 3x ploidy data set. The generation of the data set is described in Section 3.2. The error bars represent the lowest and highest values for the metric over three iterations. The Hamming and switch error rates are averaged over all broken blocks for WHP.

We found that WHP takes a conservative approach as *>* 15% of the variants were not called by WHP, even on 20x coverage per haplotype data. This is included as part of the Hamming error rate in our tests. In [1], the authors also identify that WHP tends to be extremely conservative with respect to cutting haplotype blocks and calling variants. One of the implications is that the Hamming error rate for WHP on our data set is significantly higher than that reported in their study [33]. However, there are several differences in the settings for our respective studies, which we hypothesize may contribute to that disparity. For example, there are relatively high rates of heterozygosity in our test data set (45 bp between SNPs on average), which is less the case on the reported test results for WHP [33], as they test on artificial human polyploid chromosomes created by combining different human chromosomes; it is well known that human chromosomes have much lower rates of heterozygosity than potato genomes which have variants on average *<* 50bp apart [35].

Another difference is in the simulator we used to generate long-read data for our benchmarks (PaSS [39], NanoSim[37]), which may differ in error profile than the data in their experiments.

Finally, we noticed that on polyploid genomes that have very similar haplotypes, i.e. genomes which exhibit heavy amounts of collapsing, the run time of WHP is much faster than on our simulated data set. Regardless, we believe that more testing is needed to identify the regimes in which the various types of long-read based phasing algorithms perform optimally.

## G Results on simulated data sets with different rates of heterozygosity

We include two additional tests based on the simulated data set described in the paper under the same testing conditions with the rates of heterozygosity changed from 45 bp to 90 bp and 135 bp between SNPs on average (mean). The results are shown in Figure 8 and Figure 9. At higher rates of heterozygosity, flopp has a higher likelihood of a major switch error occurring which may result in a high hamming error rate. However, the switch error rate for flopp is still better than H-PoP, showing that at on a local scale, flopp still performs well.

**Fig. 8.**
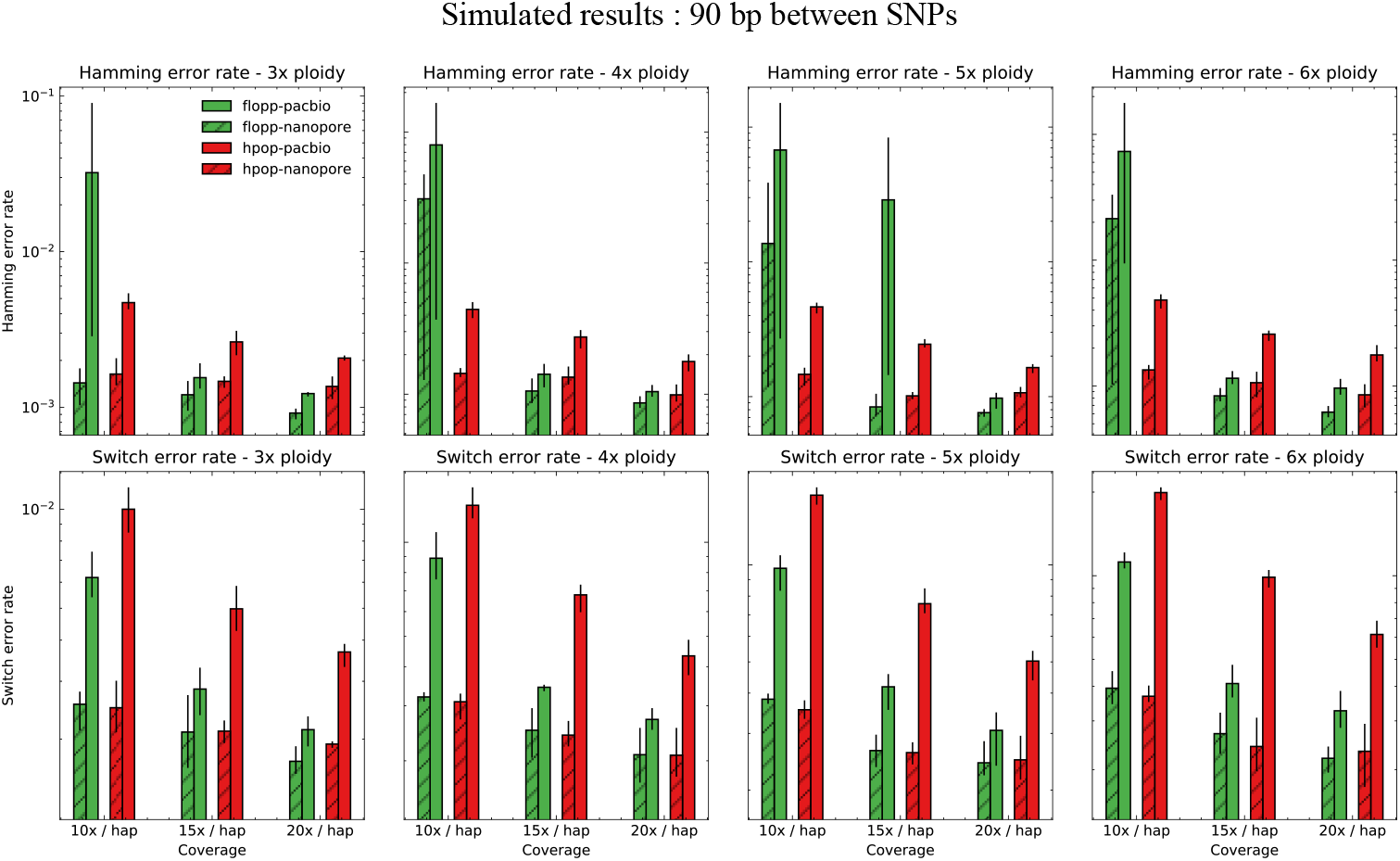
The mean switch error rate and Hamming error rate from testing on simulated data sets as described in Section 3.2 except with 90 bp between SNPs on average (instead of 45bp). The error bars represent the lowest and highest values for the metric over three iterations.

**Fig. 9.**
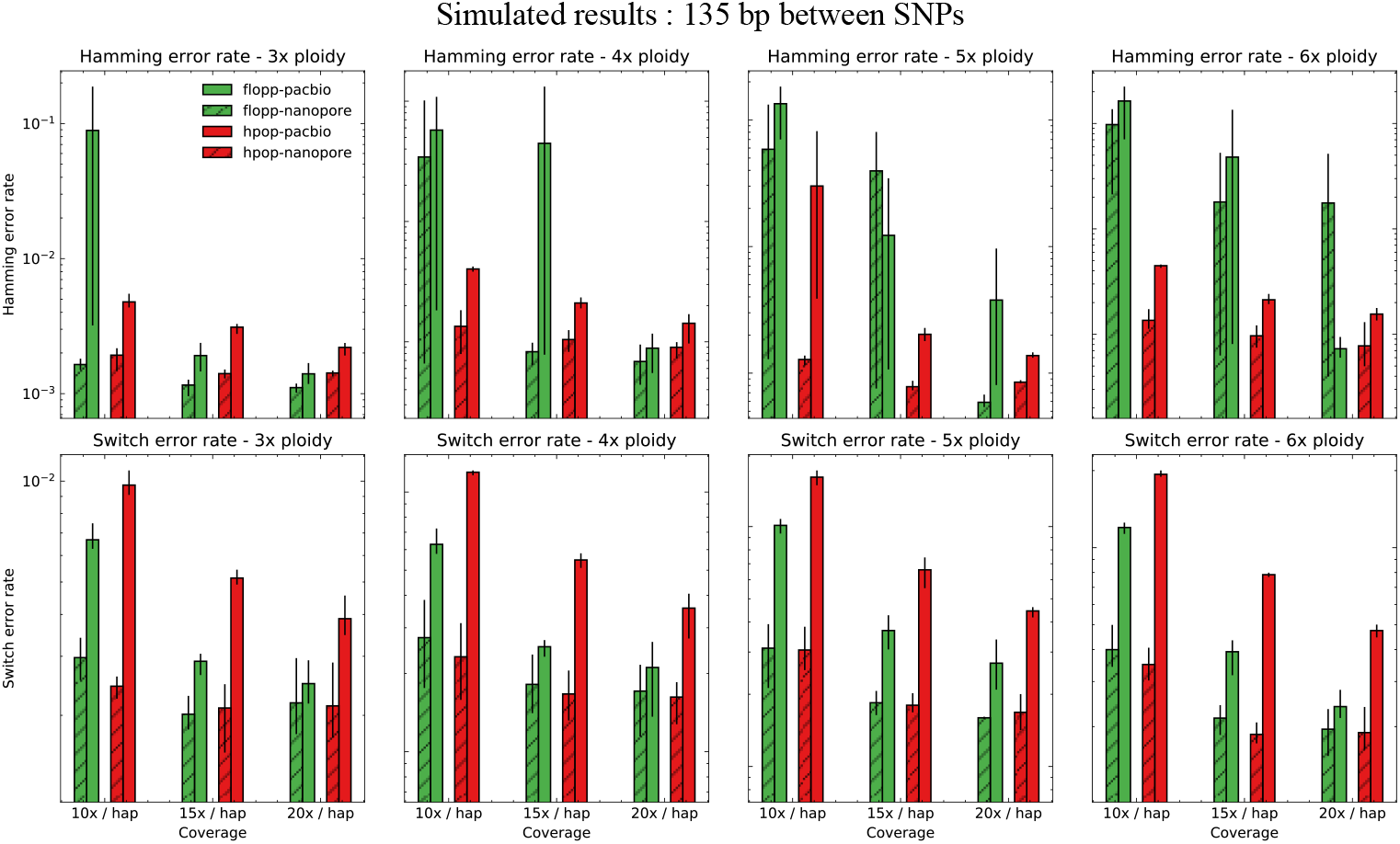
The mean switch error rate and Hamming error rate from testing on simulated data sets as described in Section 3.2 except with 135 bp between SNPs on average (instead of 45bp). The error bars represent the lowest and highest values for the metric over three iterations.

